# Magnitude estimation reveals Poisson-like noise underlying perception

**DOI:** 10.64898/2026.03.02.709083

**Authors:** Cristina Rodríguez-Arribas, Joan López-Moliner, Daniel Linares

## Abstract

Understanding how physical stimuli map onto perceptual experience is the founding goal of psychophysics. Insights into this process have primarily relied on measurements of sensitivity—how finely people tell stimuli apart—but these cannot identify the internal noise distribution underlying perception because they cannot separate it from signal amplification. Here, using visual contrast, we show that the trial-to-trial variability of subjective ratings derived from magnitude estimation—where people rate stimulus intensity with numbers—provides access to this noise. We validated this estimate by demonstrating that the analysis of this variability alongside mean responses reveals an internal representation comprising a sigmoidal response function and Poisson-like noise that quantitatively predicts sensitivity without free parameters. These predictions include the classic phenomena of the pedestal effect and Weber’s law, which we can now trace back to the response nonlinearities and signal-dependent noise. These findings establish subjective rating variability as a behavioral measure of internal noise.

## Main text

Sensory processing is inherently noisy. Even repeated presentations of the same stimulus elicit variable neural responses and variable perceptual judgments (Britten et al., 1992; Green & Swets, 1966). Since Fechner founded psychophysics, this variability has been characterized through discrimination sensitivity, which quantifies how reliably an observer can tell apart two similar stimuli, such as those differing slightly in motion strength or stimulus intensity (Britten et al., 1992; Fechner, 1860; Gescheider, 1997; Green & Swets, 1966).

Discrimination sensitivity, however, provides an incomplete picture of how a stimulus is internally represented (Green & Swets, 1966; Seriès et al., 2009; Zhou, Duong, et al., 2024). High sensitivity could arise for two fundamentally different reasons: either the sensory system strongly amplifies small stimulus changes—so that the average internal response differs markedly between similar stimuli (high signal gain)—or the internal response to a given stimulus is highly reliable from trial to trial (low internal noise).

This signal-to-noise ambiguity is built into the very definition of sensitivity under signal detection theory (Green & Swets, 1966; Seriès et al., 2009; Zhou, Duong, et al., 2024). Within this framework, the local sensitivity *D*(*x*) to discriminate a stimulus intensity *x* from *x* + Δ *x* is defined as:

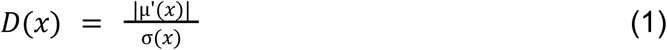

Here, the numerator represents the signal amplification, or the gain of the transducer (the function µ(*x*) mapping the physical stimulus to the mean internal response) and the denominator σ(*x*) represents the trial-by-trial internal noise. Because behavioral discrimination only measures the resulting ratio, it cannot isolate the transducer from the noise.

Resolving this underdetermination requires a method that can empirically constrain the transducer and the noise of the internal representation simultaneously. Recently, magnitude estimation (ME)— the central task of Steven’s psychophysics, in which observers provide subjective ratings by assigning numerical values to match perceived stimulus intensities (Stevens, 1957)—has been proposed to overcome this ambiguity (Zhou, Duong, et al., 2024). The proposal builds on a mathematical demonstration that mean subjective ratings can be reconciled with discrimination sensitivity, the cornerstone of Fechner’s tradition, both being linked through Equation 1. This reconciliation rests on the long-standing assumption of Steven’s psychophysics—that mean ME responses reflect the sensory transducer (Gescheider, 1997; Stevens, 1957; Werner & Mountcastle, 1965), an interpretation that has itself been long debated (Krueger, 1989).

A stronger version of this idea, however, can be put to the test. If ME captures not just the transducer through its mean, but also the internal noise through its trial-by-trial variability—a dimension of the data that has been largely overlooked (but see (Green & Luce, 1974)—then the full internal representation could be recovered from ME alone. This is not merely an additional assumption because it yields a stringent test: the internal representation derived from ME should then quantitatively predict discrimination sensitivity, with no free parameters.

We tested this hypothesis using visual contrast, a canonical attribute for studying perceptual processing (DeValois & DeValois, 1990). Because discrimination alone cannot resolve the behavioral underdetermination described above (Georgeson & Meese, 2006), models of contrast perception have typically imposed assumptions about the structure of internal noise. These assumptions range from constant noise (Boynton et al., 1999; Legge & Foley, 1980; Pelli, 1985; Pestilli et al., 2011; Wilson, 1980), to multiplicative noise (Chirimuuta & Tolhurst, 2005), mixed additive and multiplicative noise (Zhong-Lin Lu et al., 2008), and biologically motivated forms of variability (May & Solomon, 2015). Rather than imposing a noise structure, our approach seeks to estimate it from trial-by-trial ME variability.

Contrast provides a robust validation of this approach because its sensitivity function is strongly non-monotonic (Legge, 1981; Nachmias & Sansbury, 1974), a hallmark observed across sensory modalities (Arabzadeh et al., 2008; Green, 1960; Hanna et al., 1986). At very low intensities, discrimination improves with intensity (the pedestal effect) (Legge, 1981; Nachmias & Sansbury, 1974), whereas at higher intensities, it deteriorates (Weber-like behavior) (Fechner, 1860; Gescheider, 1997). By collecting ME responses across contrasts spanning these two regimes, we tested whether their full distribution can predict these complex, nonlinear behavior.

## Results

### ME reveals a sigmoidal transducer and Poisson-like noise

We presented gratings with different contrasts ranging from 0.05% to 14% and asked 11 participants to report their perceived intensity using any number of their choice (ME; Fig. 1a). We modeled the response distributions using a normal probability *N*(µ(*x*), σ(*x*) ^2^), systematically testing different functional forms for the transducer and noise (fig. S1 to S4). We found that for most participants the data were best described by a model combining a non-saturating sigmoidal transducer (Fig. 2. A and B, Eq. 2) with power-law noise including an additive component (Fig. 2C, Eq. 3, and table S1).

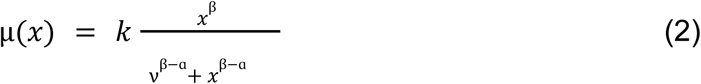

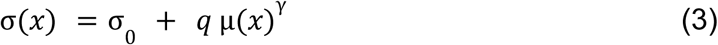

**Fig. 1.**
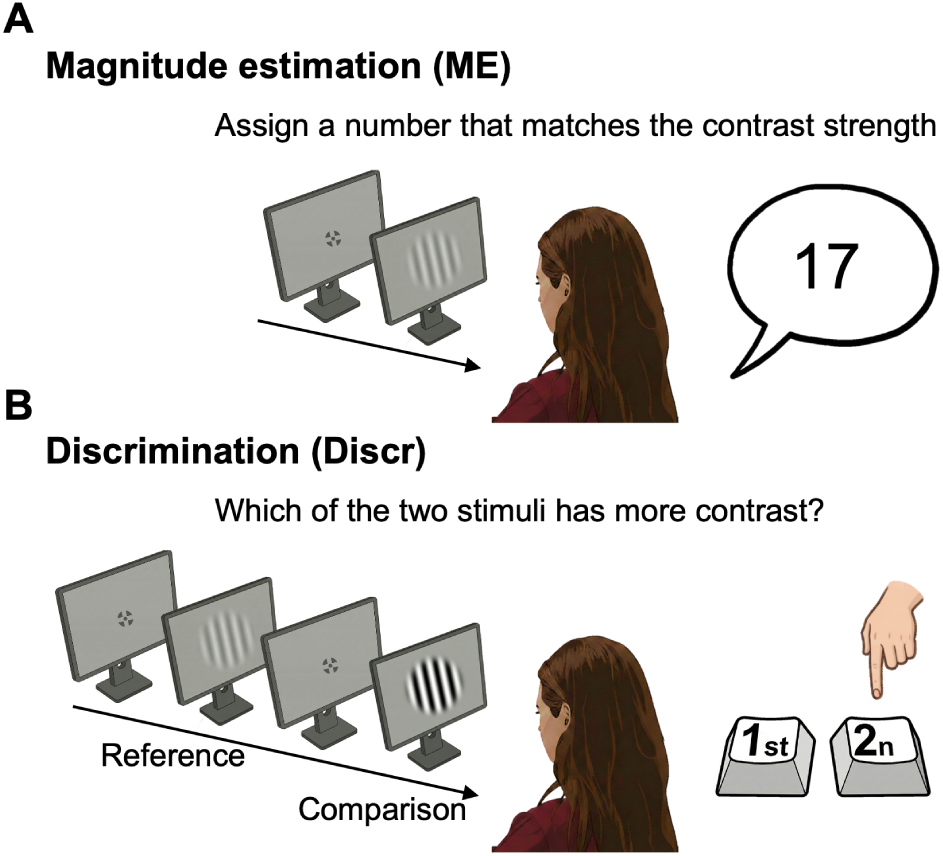
Illustration of the tasks. (**A**) In the magnitude estimation task (ME), participants rate stimuli contrasts using unconstrained verbal numerical reports. (**B**) In the discrimination task, participants compare references and comparison stimuli with both their presentation order and contrast values randomized across trials.

**Fig. 2.**
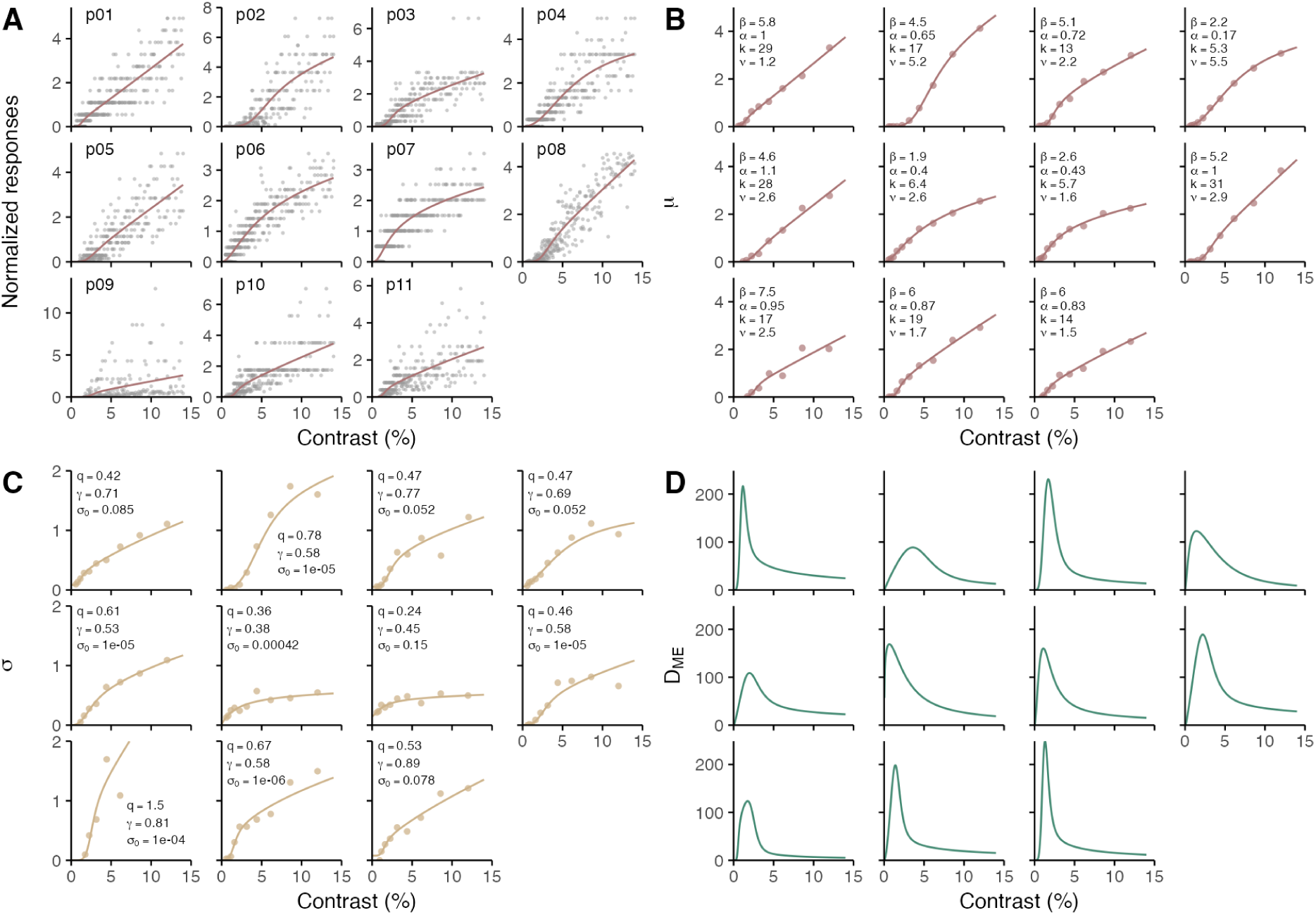
ME predicts hallmark discrimination phenomena. (**A**) ME responses (dots) normalized by each participant’s mean, plotted against contrast for each participant. Data for each participant are shown in separate sub-panels, and this participant order is maintained in panels **B** to **D**. The solid line represents the best-fitting transducer function (Eq. 2). (**B**) Mean response for each contrast bin (dots) and the corresponding best-fitting transducer function (line). Insets display the optimal parameters of the model. (**C**) Standard deviation of responses for each contrast bin (dots) plotted against contrast. The solid line represents the best-fitting noise function (Eq. 3), revealing a Poisson-like scaling of internal variability. Insets display the optimal parameters. Note that the data for participant p09 were truncated for visualization. (**D**) Discrimination sensitivity profiles predicted solely from ME data (transducer and noise) using Eq. 1. The predicted curves recover both the Weber-like behavior (decline at high contrasts) and the pedestal effect (initial rise).

At high contrasts (*x*≫*v*, where *v* is the semisaturation constant with a mean across participants of 2.7%; 95% CI: [1.7, 3.7]), the transducer exhibits a compressive nonlinearity approximating a power law µ(*x*) ∝ *x* ^a^, with a mean exponent of α = 0. 74 (95% CI: [0.54, 0.94]). This is consistent with earlier reports of compressive power-law behavior in contrast perception obtained using ME (Gottesman et al., 1981). Conversely, at low contrasts (*x*≪*v*), the transducer exhibits an expansive nonlinearity µ(*x*) ∝ *x* ^β^ with exponent of β = 4. 68 (95% CI: [3.49, 5.87]). This large exponent captures a rapid steepening of ME ratings at very low intensities. This departure from the simple global power-law behavior has been documented for many sensory attributes (Gescheider, 1997), including contrast (Gottesman et al., 1981).

Because ME recovers the transducer and the noise as separate functions we can now disentangle their respective contributions to the pedestal effect and Weber-like behavior.

The pedestal effect emerges because, despite a contrast-dependent increase in internal noise (Fig. 2C), the transducer’s expansive nonlinearity in the low-contrast regime (Fig. 2B) amplifies the signal rapidly enough to outpace it, resulting in heightened discrimination sensitivity.

To assess the origin of Weber-like behavior at higher contrasts, we substitute the internal representation (Eq. 2 and 3) into the general sensitivity equation (Eq. 1), yielding a sensitivity that scales as *D*(*x*) ∝ *x* ^a(1-y)-1^. Using the mean parameter estimates across participants (α = 0. 74, γ = 0. 64), the resulting exponent is α(1 − γ) − 1 = 0. 74 · (1 − 0. 64) − 1 = − 0. 73, which is consistent with Weber-like behavior (a canonical Weber’s law predicts an exponent of -1). This decomposition clarifies the relative contributions of the transducer and internal noise. If the noise were constant (γ = 0), the transducer’s compressive nonlinearity alone ( α = 0. 74) would predict a slope of 0. 74 · (1 − 0) − 1 = − 0. 26, which is insufficient to capture the steepness of the observed Weber-like behavior. Conversely, a linear transducer (α = 1) combined with our observed noise scaling ( γ = 0. 64) would yield a slope of 1 · (1 − 0. 64) − 1 = − 0. 64, which is much closer to the observed Weber-like behavior. This average pattern holds for most participants, though the transducer’s compressive nonlinearity also contributes to Weber-like behavior in 3 individuals (table S2). This suggests that Weber-like behavior primarily emerges from signal-dependent noise, with transducer compression generally playing a secondary role.

### Discrimination sensitivity matches predictions from ME

Having successfully reproduced the qualitative structure of these hallmark discrimination phenomena, we next tested whether the internal representation derived from ME could quantitatively predict empirical discrimination sensitivity. Using the same stimuli, we measured sensitivity in the same participants through a two-interval forced-choice task where they judged which of two successive stimuli had higher contrast (Fig. 1B). To obtain a continuous measure of sensitivity, we modeled the proportion of correct responses (Fig. 3A) using psychometric functions arising explicitly from a transducer µ(*x*) and internal noise σ(*x*) (García-Pérez & Alcalá-Quintana, 2007). Because discrimination data cannot uniquely constrain both functions simultaneously (Eq. 1), we used the functional forms that provided the best fit for the ME task (Eq. 2 and 3), but independently refit their parameters to this new dataset. Importantly, this choice was not critical, as conventional fitting procedures, calculating thresholds from independent logistic functions at each reference level, yielded similar sensitivities (fig. S5).

**Fig 3.**
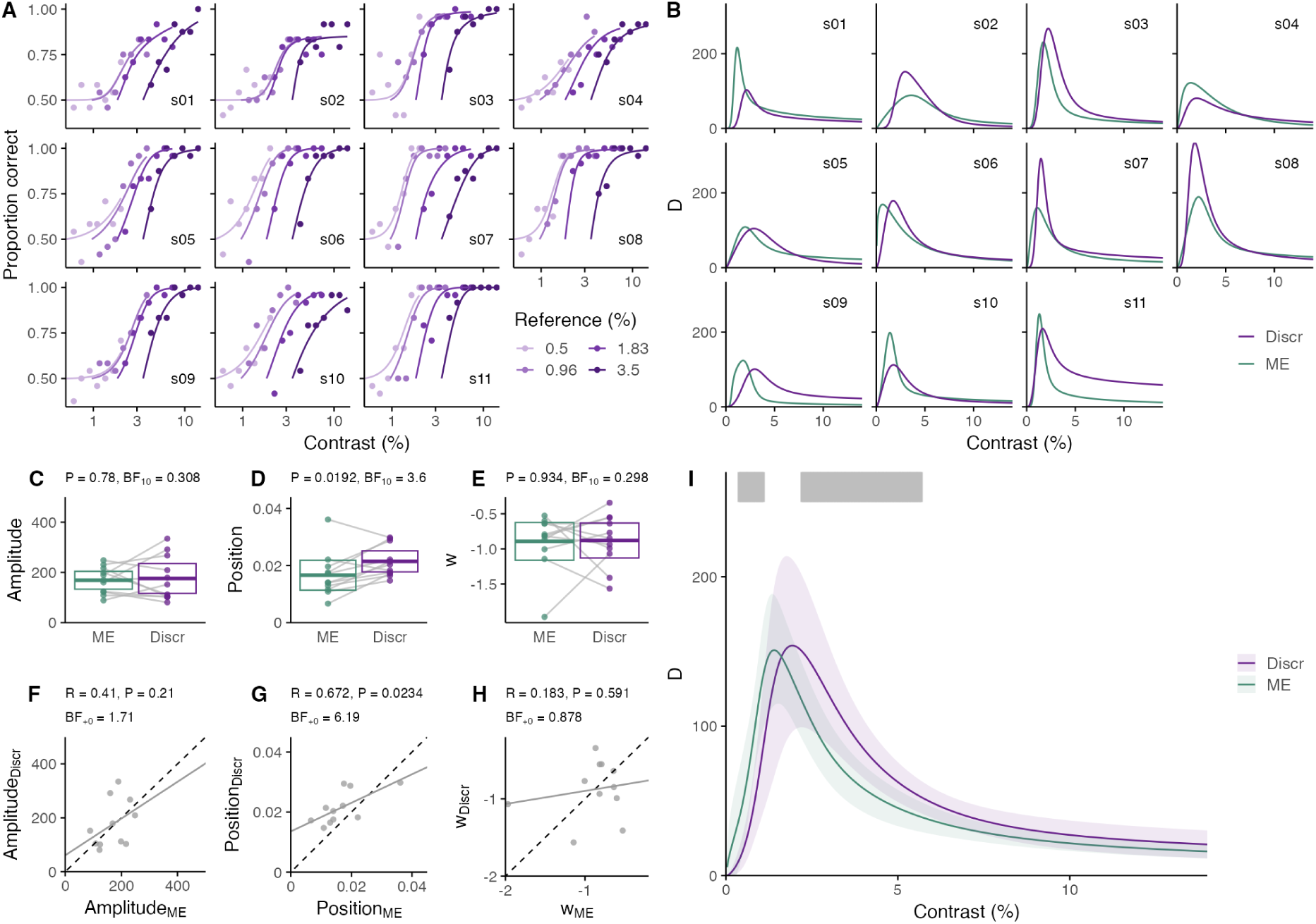
Discrimination sensitivity quantitatively matches predictions from ME. (**A**) Proportion of correct responses (dots) plotted against the log contrast of the comparison stimulus for four reference contrasts for each participant. Solid lines represent the psychometric functions that best fit this discrimination data. (**B**) Sensitivity profiles derived from the psychometric functions in (**A**) compared with predictions from the ME task. (**C** to **E**) Comparison of the peak amplitude, peak position, and Weber-like exponent between the discrimination and ME sensitivity profiles. The boxes indicate group means and t-based 95% confidence intervals. Annotations indicate P-values from two-tailed paired t-tests (P) and Bayes Factors (BF_10_) quantifying the evidence against the null hypothesis. (**F to H**) Correlations across participants for the peak amplitude, peak position, and Weber-like exponent between both tasks. Annotations indicate Pearson’s coefficient (R), two-tailed P-value (P), and one-tailed Bayes Factor correlation analysis (BF_+0_). (**I**) Mean sensitivity profile across participants for ME and discrimination tasks. Gray shaded areas indicate contrast values with significant differences between tasks (P < 0.05, two-tailed paired t-test).

As expected, the contrast discrimination task successfully reproduced Weber-like behavior and the pedestal effect in all participants (Fig. 3B). To quantify the agreement between the sensitivity predicted from ME (*D_ME_*) and the empirically observed discrimination sensitivity (*D_Discr_*), we compared three defining features of the sensitivity profiles: the peak amplitude, the peak position, and the power-law exponent characterizing the Weber-like regime.

At the group level, peak amplitudes (Fig. 3C) and Weber-like exponents (Fig. 3E) showed no systematic differences, though *D_ME_* peaked at slightly lower contrasts than *D_Discr_* (Fig. 3D), which is evident as a leftward shift in the average sensitivity profiles across participants (Fig. 3I). To assess predictive accuracy at the individual level, we evaluated whether ME could account for variations in discrimination performance (Fig. 3. F to H). We found evidence that *D_ME_* successfully captured individual differences in peak positions (Fig. 3G), but inconclusive evidence regarding peak amplitudes and Weber-like exponents (Fig. 3F and 3H).

Overall, despite the individual-level variability and the task-specific shift, the quantitative agreement of the sensitivity profiles derived from these two methodologies, which have historically been viewed as fundamentally different (Krueger, 1989), is remarkable.

## Discussion

While decades of research in the tradition of Stevens’ psychophysics have documented how the mean of subjective ratings scales with intensity (Gescheider, 1997), the variability of these responses has been largely overlooked. Here, we analyzed this variability and found that it follows a lawful, signal-dependent pattern. Importantly, when coupled with the mean response, this variability quantitatively predicts complex nonlinear discrimination phenomena—Weber-like behavior and the pedestal effect—hallmarks of Fechner’s psychophysics that have long constrained theories of sensory processing (Legge & Foley, 1980; Pardo-Vazquez et al., 2019). This validates rating variability as a behaviorally accessible measure of perceptual noise and provides direct empirical support for the foundational assumption of Stevens that the mean response reflects the sensory transducer. Ultimately, our findings show that ME effectively characterizes the core of internal representation of intensities, demonstrating that a single internal response function underlies both subjective ratings and objective discrimination. Thus, while recent work has established that Fechner’s and Stevens’ psychophysics could be mathematically compatible (Zhou et al., 2024), our results provide strong evidence that they are empirically compatible when the variability of ME is accounted for.

Our analysis of the variability of ME revealed that internal noise follows a Poisson-like noise structure. This direct measurement provides concrete empirical backing for the mathematical inferences Zhou et al. (Zhou, Duong, et al., 2024) made to reconcile existing literature. In their study, they compiled discrimination *D*(*x*) and mean subjective ratings µ(*x*) from separate studies to test their compatibility using Equation 1. They found that these two types of judgments could be reconciled by assuming a modulated Poisson noise structure 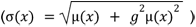 (Zhou, Duong, et al., 2024). This noise structure transitions from a Poisson regime at low intensities (σ(*x*) = µ(*x)*^0.5^), to a multiplicative regime at higher intensities ( σ(*x*) = *g*µ(*x*)). Independent support for this type of Poisson-like noise comes from findings of Pardo-Vazquez et al. (Pardo-Vazquez et al., 2019). Taking a different methodological approach, they demonstrated that discrimination sensitivity and reaction times could be jointly explained by a bounded accumulation model that explicitly assumes a Poisson noise structure (σ(*x*) = µ(*x)*^0.5^). While these previous studies suggested that Poisson-like variability underlies internal sensory representations, their conclusions relied on indirect inference. By directly measuring response variability, our results confirm the existence of this noise structure, which we then validate by demonstrating its quantitative consistency with discrimination behavior.

We have demonstrated the empirical unification of subjective and objective judgments using visual contrast. The alignment of the contrast-dependent sensitivity curves derived from ME and discrimination tasks suggests that these judgments rely on a shared internal representation. This core representation is likely situated at early processing stages, before the divergence of task-specific decision mechanisms. These later mechanisms, which include biases known to affect each task, likely account for the small displacement that we observe along the contrast axis between the sensitivity profiles derived from the two tasks (Pascucci et al., 2023; Petzschner et al., 2015; Poulton, 1968; Yeshurun et al., 2008). Despite this task-specific offset, recovering the functional form of the internal representation of contrast allows us to directly address the long-standing debate over whether the limits on contrast discrimination arise primarily from nonlinear transduction or from variable internal noise (Georgeson & Meese, 2006). Our results show that both mechanisms are necessary, but dominate sensitivity at different contrast regimes. At low contrasts, the pedestal effect emerges from the transducer’s expansive nonlinearity, which acts as an effective exponentiation threshold that amplifies signal gain sufficiently to overcome internal noise. At higher contrasts, the transducer compression plays a secondary role and Weber-like behavior arises primarily from the signal-dependent Poisson-like noise, which progressively overtakes signal gain to attenuate sensitivity.

Having recovered both the transducer and the internal noise directly from behavior, we can now ask how these two components relate to neural mechanisms—a comparison that discrimination data alone could not support due to the signal-to-noise underdetermination. Notably, the functional forms that we identify are both observed in single neurons of the visual cortex: the contrast response functions of individual units are sigmoidal (Albrecht & Hamilton, 1982), in line with normalization-based accounts of visual coding (Heeger, 1992), and their response variability grows with the mean in a slightly supra-Poisson manner (Goris et al., 2014; Tolhurst et al., 1981). How these single-cell properties map onto the population code read out for perception, however, remains unclear (Kohn et al., 2016; Panzeri et al., 2022; Renart & Machens, 2014; Zhou, Chun, et al., 2024). Our findings help constrain this relationship by suggesting that both functional forms are preserved at the macroscopic scale. For the transducer, the sigmoidal shape is evident in population-level responses measured with fMRI (Boynton et al., 1999) and EEG (Norcia et al., 2015). For the internal noise the picture is less clear, because population variability also depends on how the responses of individual neurons covary (Kohn et al., 2016; Panzeri et al., 2022; Renart & Machens, 2014; Zhou, Chun, et al., 2024); nonetheless, the signal-dependent noise that we recover is in line with recent evidence that part of cortical population variability reflects shared, gain-like fluctuations (Goris et al., 2014; Li et al., 2026).

## Materials and methods

### Participants

Eleven participants (7 males; age range 25-45 years), including the first and last authors, with normal or corrected-to-normal visual acuity participated after written informed consent was obtained. Five participants were experienced in psychophysical tasks, two of whom were familiar with discrimination tasks, while none had prior experience with ME tasks. The study was approved by the University of Barcelona Ethics Committee.

### Apparatus and Stimuli

Stimuli were generated using PsychoPy (Peirce et al., 2019) and displayed on a gamma-corrected monitor (ASUS ROG PG258Q; 1920 × 1080 pixels) viewed from a distance of 57 cm in a lit room. The stimuli consisted of vertical Gabor patches (standard deviation of the Gaussian envelope: 0.5°; spatial frequency: 1.5 cycles/°) and a fixation cross (height: 0.58°, 101.4 cd/m^2^) on which participants were instructed to maintain fixation. All stimuli were presented against a uniform grey background (180 cd/m^2^). To achieve high bit-depth resolution for contrast presentation, we implemented the noisy-bit method (Allard & Faubert, 2008).

### Experimental procedure

#### ME task

Each trial began with a fixation point displayed for 500 ms, followed by a 500 ms blank interval, and concluded with a Gabor patch presented for 100 ms. Then, participants provided a verbal numerical estimate of the perceived contrast using an unrestricted numerical scale (allowing both integers and decimals), which the experimenter recorded manually. The Gabor patches were presented at 200 contrast levels, logarithmically spaced between 0.5% and 14%. Each contrast level was presented once per session in a random order across two separate sessions, resulting in a total of 400 trials per participant. Prior to these sessions, participants completed a practice block of 30 trials to stabilize their use of the internal response scale.

#### Discrimination task

Before conducting the ME task, participants performed a two-interval forced-choice discrimination task. Each trial began with the presentation of a fixation point for 500 ms, followed by a 500 ms blank interval. Then, the reference and comparison stimuli were presented sequentially in random order for 100 ms each, separated by an inter-stimulus interval of 1000 ms. Participants indicated which interval contained the stimulus with higher contrast using a keyboard response.

Four reference contrast levels (0.5%, 0.96%, 1.83%, and 3.5%) were each paired with seven logarithmically spaced comparison contrasts. The comparison ranges were 0.61–2% for the 0.5% reference, 1.17–3.84% for the 0.96% reference, 2.23–7.32% for the 1.83% reference, and 4.27–14% for the 3.5% reference. For every reference-comparison pair, there were eight repetitions, consisting of two presentation orders (reference or comparison first) crossed with four spatial phases (0, π/2, π, 3/2π). Participants completed three blocks of trials, resulting in a total of 672 trials (4 references x 7 comparisons x 8 repetitions x 3 blocks). The trial order was fully randomized within each block.

## Data Analysis

Analysis and figures were conducted using the R statistical computing environment (R Core Team, 2025).

### ME task

To ensure data quality prior to curve fitting, we preprocessed the data to address scale usage, outliers, and zero-inflation. First, to assess the stability of scale usage, we estimated the standard deviation of responses across consecutive trial bins. This analysis revealed unusually large response variability during the initial trials of three participants (p03, p09 and p11), leading to the exclusion of the first 10% of their trials. Next, outliers were identified by fitting a LOESS curve to the raw data. Responses falling outside ± 3 standard deviations from the fitted curve were excluded, resulting in the removal of only 9 trials across the entire dataset. Finally, during preliminary curve fitting, we observed poor model fits at high contrast levels for three participants (p05, p10, p11). This was driven by a heavy concentration of zero responses at low contrasts, which disproportionately biased the maximum likelihood estimation. To restore the balance of the likelihood function, we randomly downsampled the zero responses for these specific participants, removing 21%, 18%, and 35% of their total trials, respectively.

To facilitate comparisons across participants, the ME responses were normalized by dividing each participant’s trial responses by their own overall mean. We modeled these normalized responses *r* using a normal distribution, *N*(µ(*x*), σ(*x*)), and fitted its parameters by maximizing the log-likelihood function:

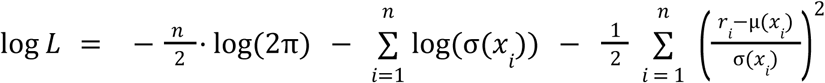

where *n* is the number of responses, µ(*x*) represents the transducer and σ(*x*) the internal noise. By considering different forms of the transducer and the noise, we evaluated a comprehensive set of candidate models (figs. S1 to S4). We performed the optimization using the R package DEoptim (Ardia et al., 2022), which performs global optimization over bounded parameter spaces. Model selection (tables S1) was performed using the Akaike Information Criterion (Burnham & Anderson, 2013).

### Discrimination task

We modeled the proportion of correct responses in the two-interval forced-choice task within a signal detection theory framework (García-Pérez & Alcalá-Quintana, 2007). In this approach, psychometric functions for all reference levels are fitted simultaneously by assuming an underlying transducer function, µ(*x*), and an internal noise function, σ(*x*). However, discrimination data alone cannot uniquely identify either the underlying functional forms or their specific parameters (Eq. 1). For the functional forms, adopting the best model established by our magnitude estimation data (Eq. 2 and 3) emerges as the natural choice. As for the parameters, we left them free to vary during the fitting process, but their values remain underdetermined by the signal-to-noise constraint. Consequently, these parameters serve exclusively as a computational tool to extract the continuous empirical sensitivity profile and should not be considered reliable for direct comparison with those obtained from magnitude estimation.

Within this framework, the perceptual responses elicited by the reference and the comparison stimuli were modeled as *R* ∼ *N*(µ(*x*), σ(*x*) ^2^) and *R_c_* ∼ *N*(µ(*x* + Δ*x*), σ(*x* + Δ*x*) ^2^), respectively, where *x* represents the contrast of the reference stimulus and *x* + Δ*x* represents the contrast of the comparison stimulus. Decisions were assumed to be based on the difference between these two independent perceptual responses, *R_dif_* = *R_c_* − *R_r_* , which is distributed as *N*(µ(*x* + Δ*x*) − µ(*x*) , σ(*x*) ^2^+ σ(*x* + Δ*x*) ^2^)

The expected proportion of correct responses corresponds to the probability that this difference is positive, indicating that the comparison stimulus elicited a stronger response than the reference. Evaluated across values of Δ*x*, this probability defines the predicted psychometric functions:

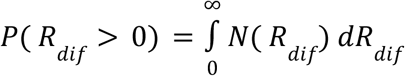

To estimate the best-fitting parameters for the chosen functional forms, we used maximum likelihood estimation, maximizing the binomial log-likelihood across all reference levels simultaneously using the R package nloptr (Johnson, 2008). Finally, we calculated sensitivity using Eq. 1.

We used this joint-fitting approach because it naturally provides a continuous measure of sensitivity across all contrast levels. Importantly, fitting independent psychometric functions for each reference level and estimating sensitivity as the inverse of the threshold yielded comparable results (fig. S5).

### Statistical analysis

We conducted mean comparisons using paired two-tailed Student’s t-tests and Bayes factor analyses with standard Cauchy priors (scale = 0.707). We assessed the strength and direction of linear associations using Pearson correlation coefficients (R) and one-tailed Bayesian correlation analyses (BF_+0_) to test for positive associations. In accordance with recent guidelines (Keysers et al., 2020), we employed directional priors for hypotheses with a predicted direction to mitigate the bias against the null hypothesis that can arise from two-tailed Bayesian tests.

## Acknowledgments

We thank Cristina de la Malla and Albert Compte for comments on the manuscript.

## Funding

Ministry of Science and Innovation fellowship PRE2021-099277 (CRA) PCI2024-155051-2 grant MICIU/AEI/10.13039/501100011 (JLM) PID2023-151752NB-I00 grant MICIU/AEI/10.13039/501100011 (DL)

## Data, code, and materials availability

All data and the code to reproduce the statistical analysis and create the figures is available on GitHub: https://github.com/viscalab/me_noise

## Supplementary Materials

Figs. S1 to S5 Table S1 to S2

---

**Fig. S1.**
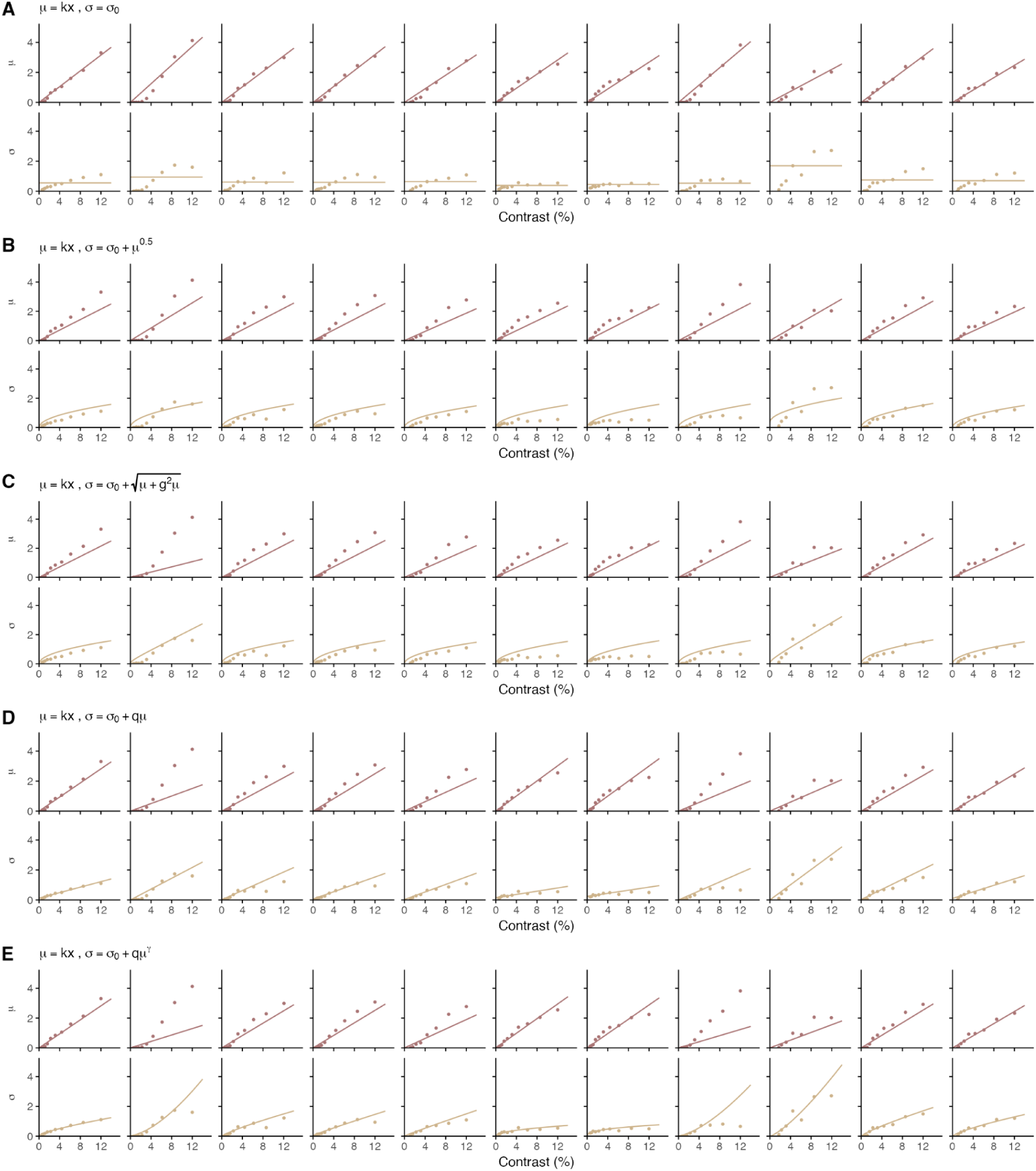
Model fits for a linear transducer coupled with alternative noise functions. All models assume a linear transducer but vary in their internal noise assumptions: (**A**) additive, (**B**) Poisson, (**C**) modulated Poisson, (**D**) multiplicative, and (**E**) power. Each column represents an individual participant (ordered from left to right as p01 to p11, matching the labels in the main text). Within each panel, the top row shows the mean response (μ) and the bottom row shows the response standard deviation (σ) as a function of stimulus contrast. Dots indicate the empirical estimates for each contrast bin, while solid lines represent the corresponding model fits. To prevent parameters from increasing indefinitely during the fitting process, we applied an upper bound of 2 to parameter g for participant p02 in the model in (**C**), and to parameter q for participants p02 and p08 in the model in (**E**). In all other cases, the optimized parameters did not hit their boundary constraints.

**Fig. S2.**
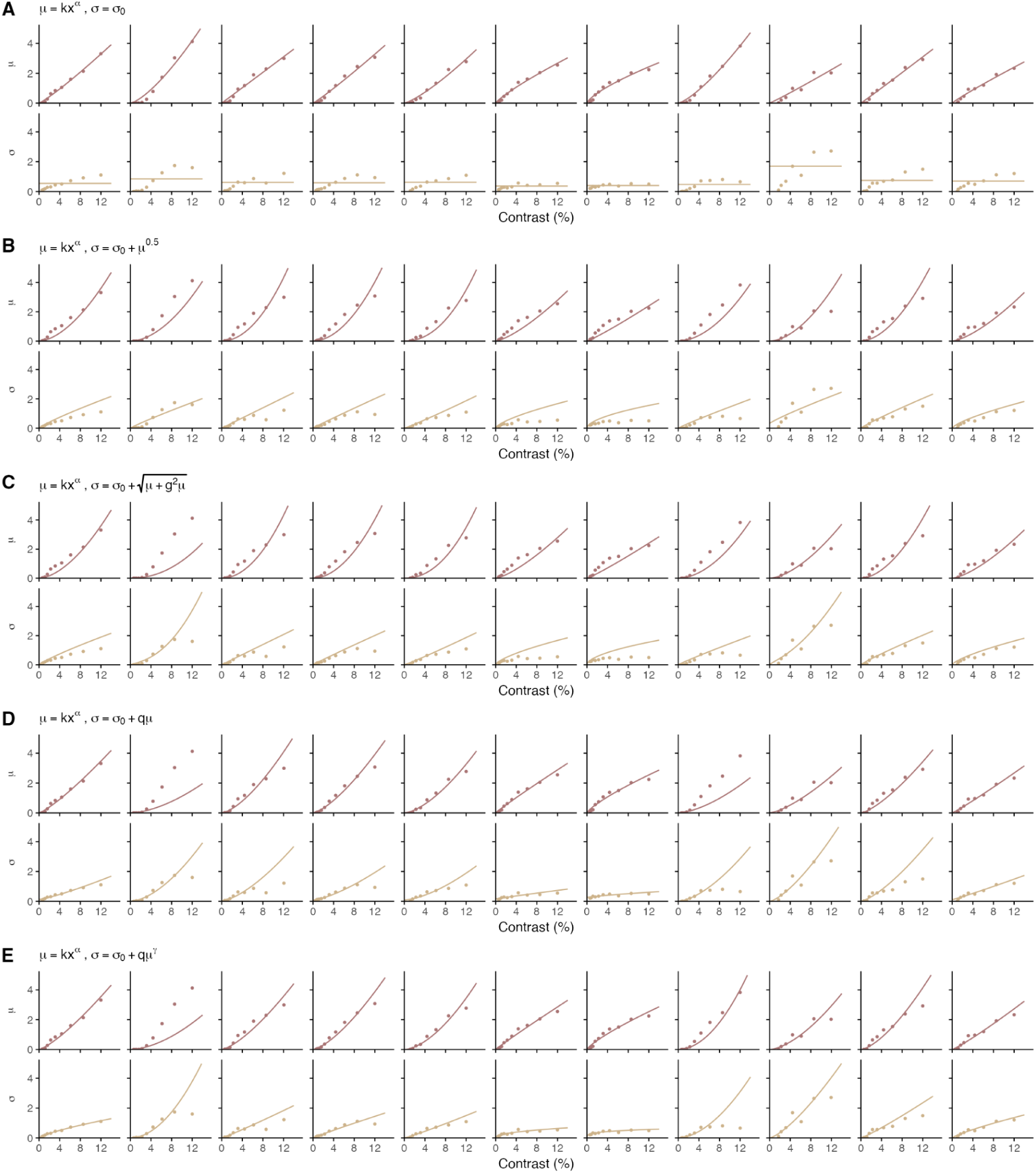
Model fits for a power law transducer coupled with alternative noise functions. Figure layout and conventions are identical to those in fig. S1 . The panels compare fits for the same five internal noise models ((**A**) additive, (**B**) Poisson, (**C**) modulated Poisson, (**D**) multiplicative, and (**E**) power noise), but assuming a power-law transducer. To prevent parameter k from increasing indefinitely during the fitting process, we applied an upper bound—defined by the mean of the remaining participants—to parameter k for participants p02 and p08 in the models in (**B**) through **(E**). As a result of this restriction, for participant p02, the optimized value of parameter g for the model in (**C**) and parameter q for the models in (**D**) and (**E**) reached their upper bound of 2. In all other cases, the optimized parameters did not reach their boundary constraints.

**Fig. S3.**
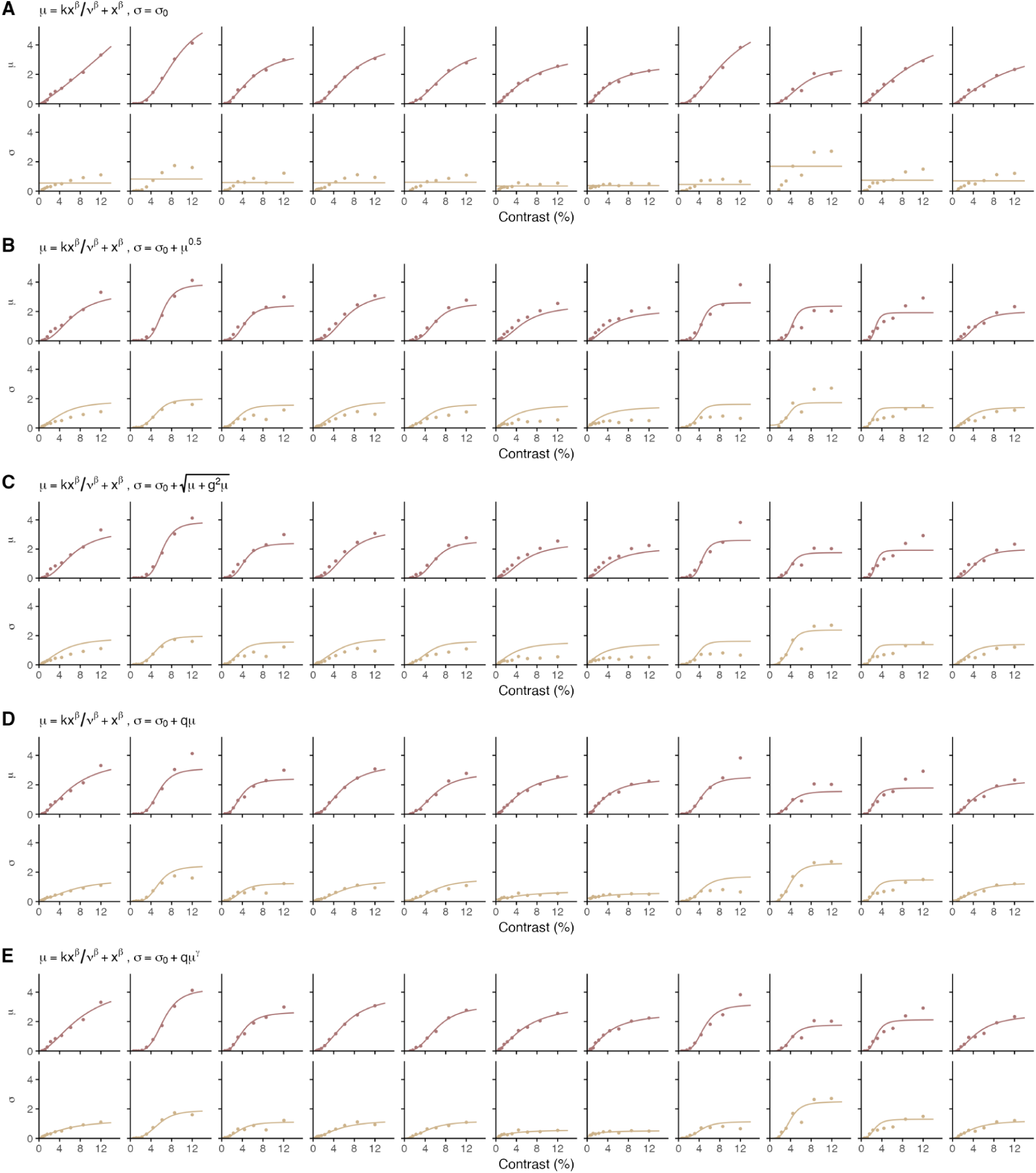
Model fits for a saturating sigmoidal transducer coupled with alternative noise functions. Figure layout and conventions are identical to fig. S1. The panels compare fits for the same five internal noise models ((**A**) additive, (**B**) Poisson, (**C**) modulated Poisson, (**D**) multiplicative, and (**E**) power noise), but assuming a saturating sigmoidal transducer. The optimized parameters did not reach their boundary constraints.

**Fig. S4.**
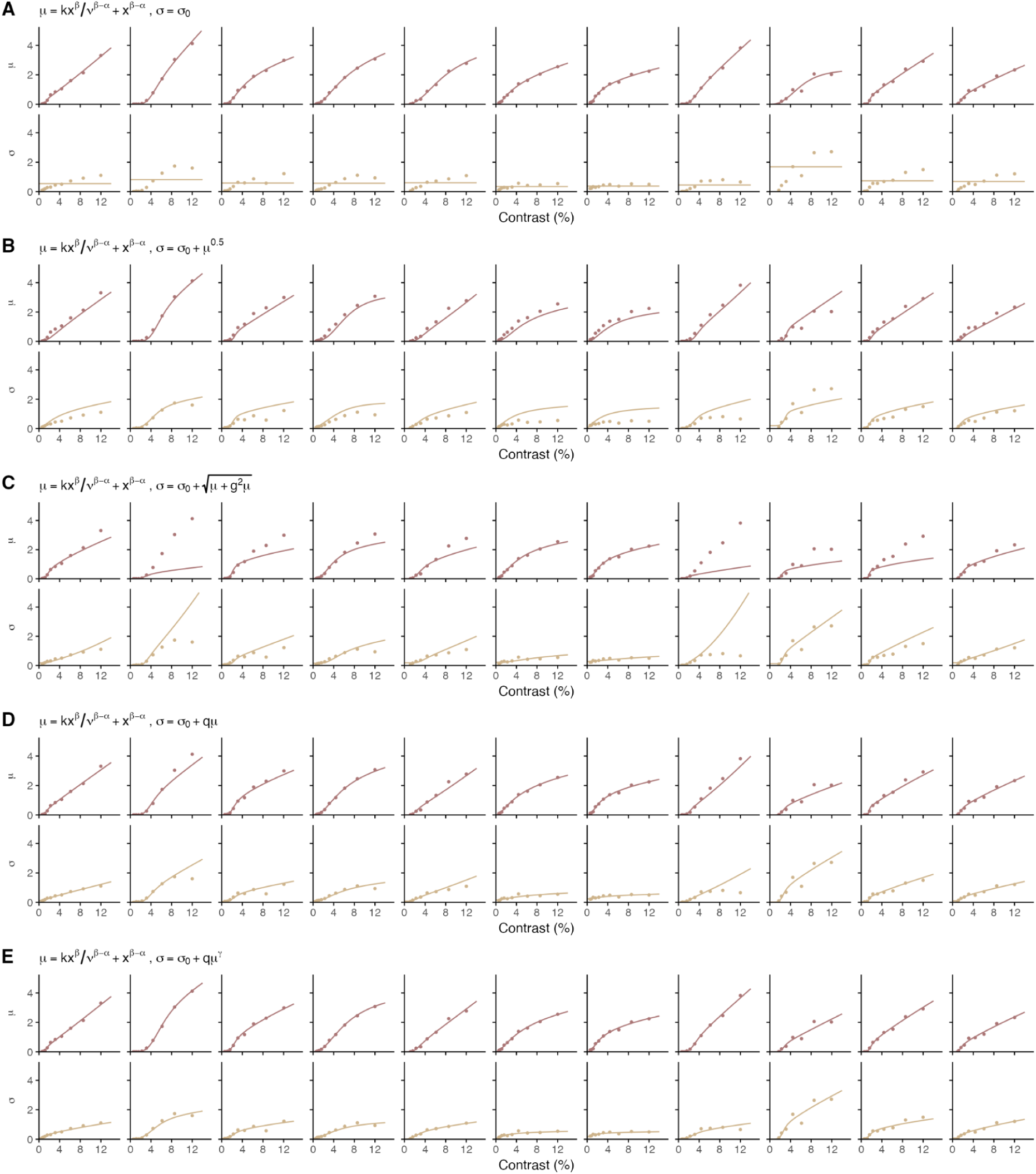
Model fits for a non-saturating sigmoidal transducer coupled with alternative noise functions. Figure layout and conventions are identical to Supplementary Figure 1. The panels compare fits for the same five internal noise models ((**A**) additive, (**B**) Poisson, (**C**) modulated Poisson, (**D**) multiplicative, and (**E**) power noise), but here all models assume a non-saturating sigmoidal transducer. The optimized parameters did not reach their boundary constraints.

**Fig. S5.**
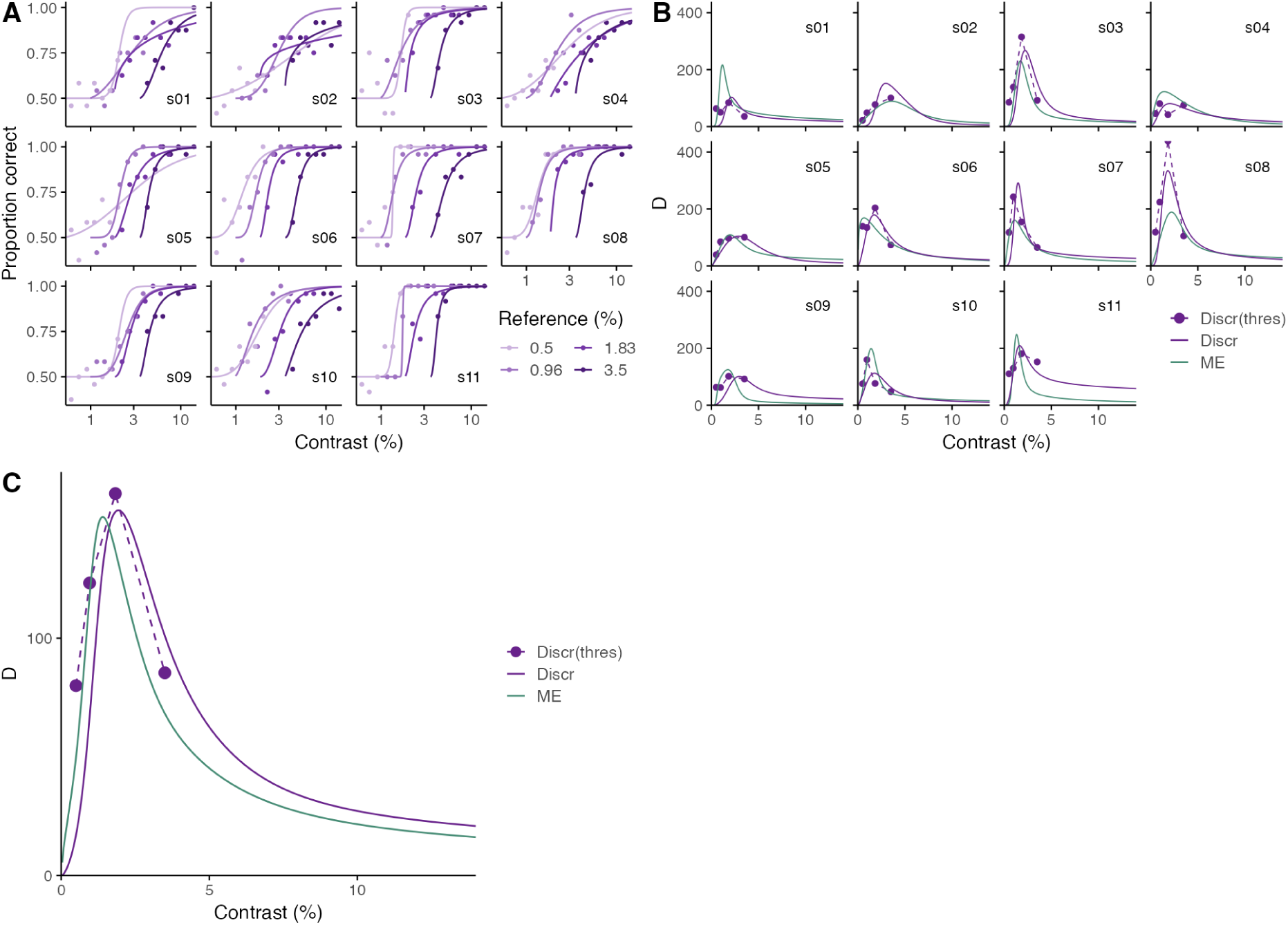
Discrimination sensitivity estimated from thresholds of independently fitted psychometric functions. (**A**) Proportion of correct responses (dots) plotted against the log contrast of the comparison stimulus for four reference contrasts for each participant. Solid lines represent the best-fitting psychometric functions fitted independently to each reference level and participant using a logistic function. (**B**) Sensitivity computed as the inverse of the threshold (Δx^−1^)for the four reference levels (dots), derived from the independent fits in (A). For comparison, the sensitivity profiles (lines) from Fig. 3B are replotted. Thresholds were defined at the 76% performance level, which corresponds to a detectability index d’ of 1 in a two-alternative forced-choice task, where the proportion of correct responses is given by 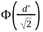 and Φ is the standard cumulative normal function (Kingdom & Prins, 2016; Seriès et al., 2009). Under this condition, given the general relationship 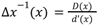 , sensitivity estimated as the inverse of the threshold coincides with the sensitivity metric defined in Equation 1 (Seriès et al., 2009; Zhou, Duong, et al., 2024). (**C**) Mean sensitivities across participants for the four reference levels (dots) plotted together with the mean sensitivity profiles (lines) from Fig. 3I.

**Table S1.**
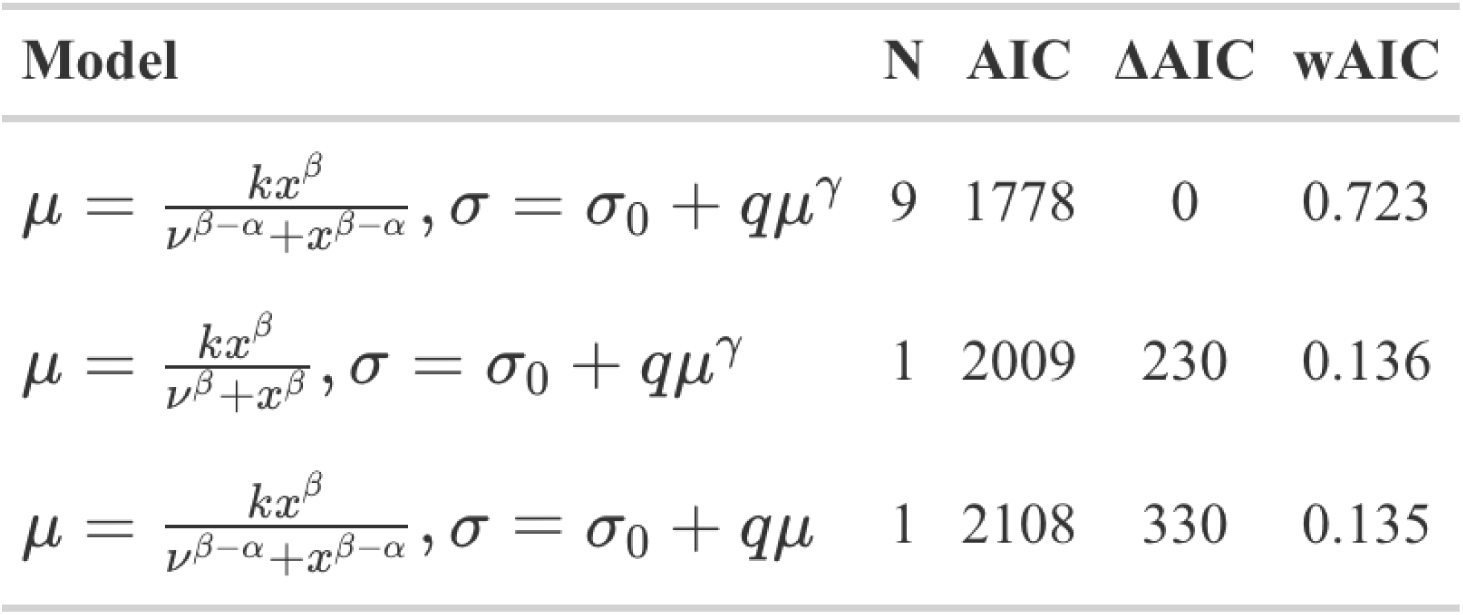
Model selection summary for the candidate models from figs. S1-S4 that were the best fit for at least one participant. The best-fitting model for each participant is identified by the lowest Akaike Information Criterion (AIC) score. N represents the number of participants for whom a given model provided the best fit. To evaluate performance at the group level, the reported AIC corresponds to the sum of individual AIC scores across all participants. ΔAIC represents the difference between a given model’s summed AIC and the summed AIC of the best-performing model in the candidate set (where ΔAIC = 0). Finally, wAIC indicates the Akaike weights averaged across all participants, reflecting the mean probability of each model being the best fit among the entire set of candidate models in figs. S1 to S4.

**Table S2.**
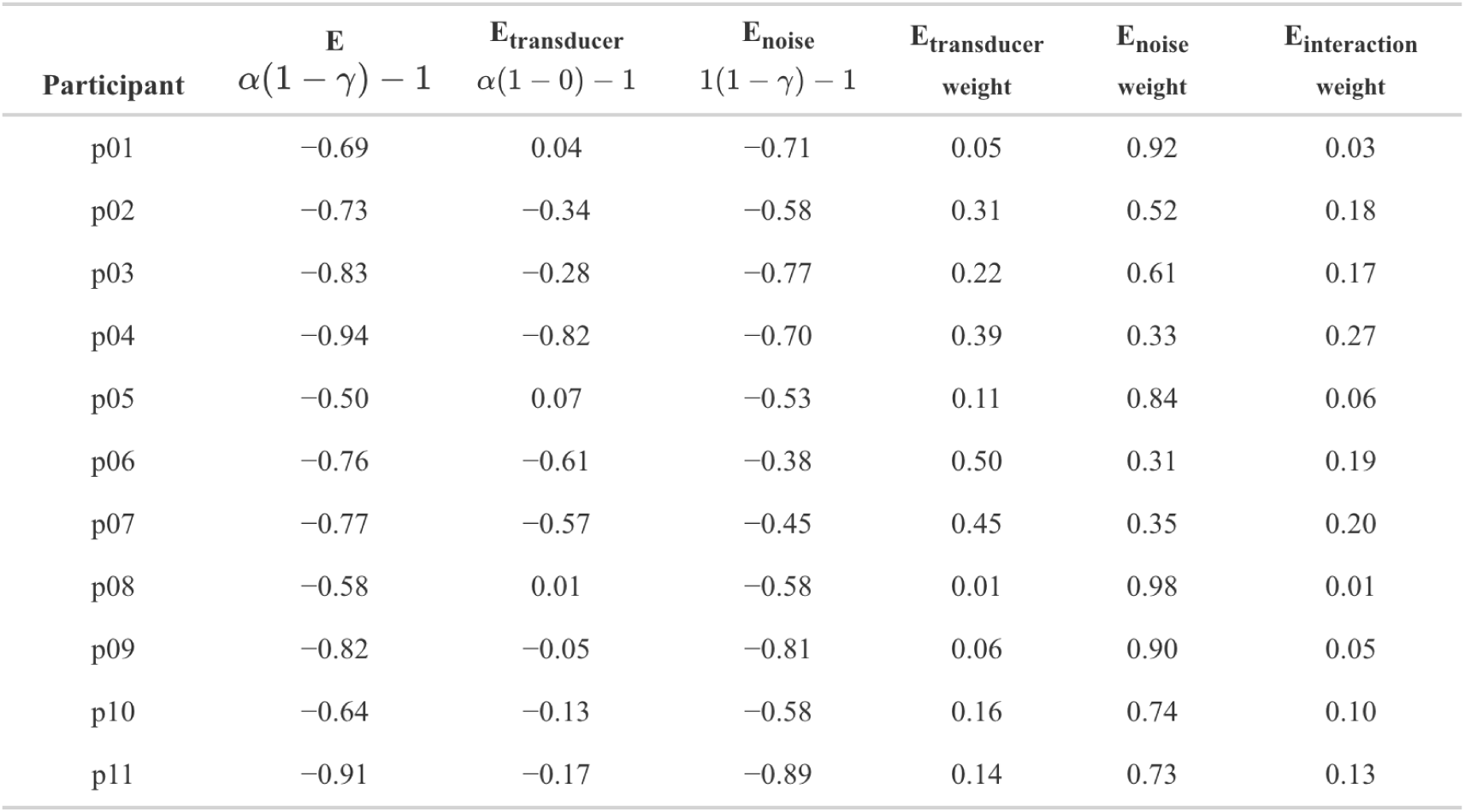
Disentangling the contributions of signal transduction and internal noise to Weber-like behavior across individual participants using ME. The first column identifies the participant. The second column indicates the asymptotic exponent of sensitivity decay at high contrasts, D(x) ∝ x ^a(1-y)-1^. The third and fourth columns isolate the specific mechanistic contributions of the transducer and the internal noise by showing the predicted decay exponent assuming constant internal noise (γ = 0) and a linear transducer (α = 1), respectively. To precisely quantify how much the transducer and the signal-dependent noise contribute to the Weber-like behavior, we decomposed the decay exponent into main effects and their interaction by centering the parameters around a baseline of 1 and 0, respectively. The exponent is, thus, rewritten as E_transduce_ + E_noise_ + E_interaction_ , where E_transducer_ = α − 1, E_noise_ =− γ and E_interaction_ =− γ(α − 1). The weight of each component is calculated by dividing its term by the total exponent (e.g., |E _transducer_| / ( |E_transducer_| + |E_noise_| + |E_interaction_ |)). The fifth, sixth, and seventh columns present these weights for the transducer, signal-dependent noise, and their interaction, respectively. The weight of E_noise_ suggests that signal-dependent noise, and not transducer compression, is the primary source of Weber-like behavior. This observation holds true for the majority of the sample; however, in three participants (p04, p06, p07), the compressive nonlinearity of the transducer also played a significant role.

| Participant | $E$<br>$\alpha(1 - \gamma) - 1$ | $E_{\text{transducer}}$<br>$\alpha(1 - 0) - 1$ | $E_{\text{noise}}$<br>$1(1 - \gamma) - 1$ | $E_{\text{transducer}}$<br>weight | $E_{\text{noise}}$<br>weight | $E_{\text{interaction}}$<br>weight |
| --- | --- | --- | --- | --- | --- | --- |
| p01 | -0.69 | 0.04 | -0.71 | 0.05 | 0.92 | 0.03 |
| p02 | -0.73 | -0.34 | -0.58 | 0.31 | 0.52 | 0.18 |
| p03 | -0.83 | -0.28 | -0.77 | 0.22 | 0.61 | 0.17 |
| p04 | -0.94 | -0.82 | -0.70 | 0.39 | 0.33 | 0.27 |
| p05 | -0.50 | 0.07 | -0.53 | 0.11 | 0.84 | 0.06 |
| p06 | -0.76 | -0.61 | -0.38 | 0.50 | 0.31 | 0.19 |
| p07 | -0.77 | -0.57 | -0.45 | 0.45 | 0.35 | 0.20 |
| p08 | -0.58 | 0.01 | -0.58 | 0.01 | 0.98 | 0.01 |
| p09 | -0.82 | -0.05 | -0.81 | 0.06 | 0.90 | 0.05 |
| p10 | -0.64 | -0.13 | -0.58 | 0.16 | 0.74 | 0.10 |
| p11 | -0.91 | -0.17 | -0.89 | 0.14 | 0.73 | 0.13 |

## Notes

https://github.com/viscalab/me_noise

### Competing Interest Statement

The authors have declared no competing interest.

### Summary of Updates

Title updated ("Subjective ratings" replaced by "Magnitude estimation"); Funding information updated; Minor text edits throughout for clarity; no changes to data or analyses.

